# Attention modulates hippocampal sharp-wave ripples in humans

**DOI:** 10.64898/2025.12.02.691881

**Authors:** Michał Domagała, Leila Chaieb, Charles E. Schroeder, Rainer Surges, Florian Mormann, Juergen Fell, Marcin Leszczynski

**Affiliations:** Cognitive Science Center, Jagiellonian University, Kraków; Department of Epileptology, University Hospital Bonn; Department of Psychiatry, Columbia University College of Physicians and Surgeons, New York City; Translational Neuroscience Lab Division, Center for Biomedical Imaging and Neuromodulation, Nathan Kline Institute, Orangeburg

## Abstract

Hippocampal sharp-wave ripples (SWRs) are high-frequency events critical for memory consolidation, typically studied during sleep and quiet wakefulness. Emerging evidence suggests that SWRs also occur during active behavior, yet their role in awake cognition remains unclear. Here, we demonstrate that changes in sustained attention modulate both the occurrence of SWRs and their temporal alignment to ongoing hippocampal oscillations. Using intracranial EEG recordings from epilepsy patients performing a sustained attention to response task (SART), we first identified attentional states based on behavioral variability and subjective reports. Next, we observed that SWRs were more frequent during high-attention periods, despite the absence of memory demands. Importantly, SWRs during low-attention periods showed higher phase synchrony with low-frequency oscillations (theta to lower beta) indicating increased endogenous coordination of SWRs under low attentional focus. These findings show that SWRs are dynamically regulated by attentional engagement, supporting a broader role for ripples in active cognitive processing.

## Introduction

Hippocampal sharp-wave ripples (SWR) can been detected as large amplitude deflections (sharp-waves) in the field potential with a brief oscillatory ripple superimposed. SWRs are observed across different species including humans (1–4), non-human primates (5–7) and rodents (8). They are typically found in the hippocampus but have also been detected outside of the medial temporal lobe, including in the primary visual cortex (9). During SWRs, hippocampal cells that fire together in the formation of an initial experience, re-activate (also termed “replay”), re-exposing hippocampal circuitry to information acquired during preceding active behavior (10–11). In rodents, SWRs are most frequently observed during slow-wave sleep (12), quiescent waking and pauses in exploration (8, 10–15). This, in turn, has led to theories of its contribution to memory and neuroplasticity during inactive states (16–19).

However, SWRs can also occur during awake states (6–8,14,20). Termed “exploratory” SWRs, these have been observed in human (2, 20) and non-human primates (6,7) during active visual exploration. For example, in a visual search task SWR rate increases with repeated presentation of scenes (7). In addition, analyses of oculomotor behavior show that SWRs are enhanced prior to gaze direction towards a familiar target location. Similarly, SWR rate during viewing (i.e., encoding) of pictures is higher for subsequently remembered images, suggesting a role of SWRs in attentive processing of sensory information (2). The fact that attended elements of a scene are better remembered (21), suggests that an increased rate of hippocampal SWRs during encoding reflects an increase in focused attention that in turn, benefits memory encoding. A number of prior studies draw links between attention, memory and cortical–hippocampal interactions (22,23): (i) Attention stabilizes representations in the hippocampus separating neural patterns elicited by sensory stimuli under distinct attentional states (21) (ii) stimuli following saccades (and associated attention shifts yoked to saccades) elicit stronger hippocampal neural responses, including activity within the SWR frequency range (24) and (iii) cholinergic stimulation influences both the rate of hippocampal SWRs (25,26) and attention (27).

Despite these advancements, it remains unclear whether attention influences hippocampal SWR activity. The goal of this study was to investigate whether fluctuations in sustained attention modulate probability of hippocampal SWR occurrence. Our analysis focused on the hippocampus as it is integral to the influence of attention on neural dynamics (21,22,24,28,29), and it is a prime generator of SWRs (8). We obtained intracranial EEG recordings from the hippocampi of presurgical epilepsy patients who had performed a variant of the sustained attention to response task (SART). By analyzing variations in reaction times, we identified states of increased and reduced focal attention (i.e., “online” and “offline” attentional states, keyed to variability in the reaction time distribution) during the performance of the SART task. We then determined the rate of SWRs during these different attentional states. Our findings indicate that attention does indeed modulate the rate of hippocampal awake SWRs, with higher rates observed during intervals of increased focal attention (i.e., online state). Consistently, we also observed elevated SWR rates during focal attention, as determined by subjective thought probe assessments to aid in definition of attentional state. We also found that SWRs coupled differently to local oscillatory activity depending on the attentional state, exhibiting stronger phase-locking to low-frequency activity during periods of reduced attention, suggesting an enhanced endogenous coordination of SWRs under low-attention conditions. These findings support a model in which focal attention facilitates encoding of sensory information by increasing the rate of “exploratory” hippocampal SWRs, whereas reduced attention modulates the timing of SWRs in coordination with ongoing hippocampal oscillations.

## Results

### Attention modulates the rate of awake hippocampal SWRs

We recorded intracranial field potentials from depth electrodes implanted in the hippocampus while participants performed a variant of the sustained attention to response task (SART; Fig. 1A upper). In this task, digits 1–9 appeared every 2 seconds, and participants were asked to respond to every digit except “3,” emphasizing both speed and accuracy. Periodically, thought probes (TP) introspectively assessed participants’ momentary attentional state (“focused” vs. “mind wandering”) at an inter-probe interval of 20-30 seconds. Importantly, the task contained to no explicit memory component.

**Figure 1:**
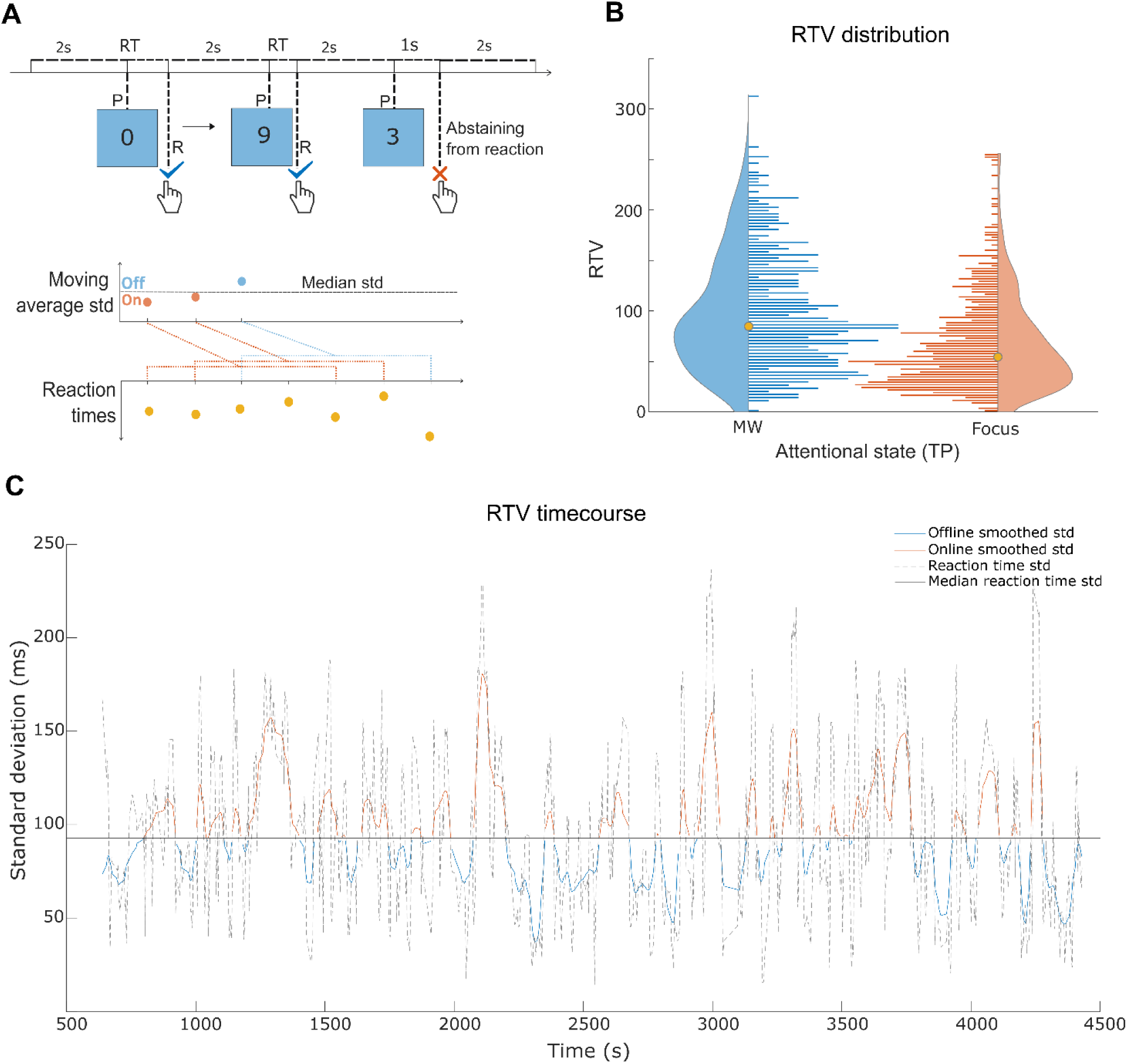
Overview of the SART task and behavioral analyses. **(A)** An overview of the SART task and Reaction Time Variability (RTV) computation. **Upper part** represents a SART scheme. Participants were presented with a random sequence of digits ranging from 0 to 9 with 2 s inter-digit interval. They were instructed to press the space bar in all cases except when the number ‘3’ was presented, in which case they had to abstain from a response (speed and accuracy were emphasized). After pressing the button, the next digit was displayed after 2 s. If no response was made, the procedure re-commenced after 1 s (plus a 2 s interval period). “P” indicates presentation of a digit, “R” corresponds to participant’s response. **Lower part** represents a moving standard deviation (std) method used for computing RTV time series which was calculated using a sliding window of 5 measurements (participant responses in this case), moving one step at a time. To estimate variability in reaction times we calculated the standard deviation across these 5 measurements. The RTV time series was then split, categorizing observations exceeding the median as offline states and those below as online states. This measure reflects an established relationship between variance in reaction time responses and sustained attention (see Methods). **(B)** Distribution of RTV preceding thought probe reports of mind wandering (MW; blue) and being focused (orange) across all participants (n = 928 thought probes across N = 12 patients). The medians are marked by large orange dots. The clouds and lines represent a distribution (cloud) and histograms (lines, number of bins = 100) of RTV in MW (blue) and Focused (orange). **(C)** An example of a single participant RTV time course. Raw RTV (dashed line) was smoothed (here with Gaussian kernel σ = 20) and binarized into offline (orange) and online (blue) attentional states.

To quantify attentional fluctuations, we computed reaction time variability (RTV) using a rolling standard deviation of five consecutive responses, with the window advancing by one response at a time (Fig. 1A/C lower; see Methods for details). Following established procedures (see Methods), periods with RTV below the median were classified as “online” (high attention), and those above the median as “offline” (low attention). Participants experienced a comparable number of online and offline states (*mean number of intervals: online* = 58.77, *offline* = 58.81, *z* = -0.49, *p* = 0.62*, N* = 17, Wilcoxon signed-rank test). Offline intervals were marginal longer compared to online intervals (*online* = 17.33 s; *offline* = 19.54 s; *p* = 0.0552, *z* = 1.917, *N* = 17). We additionally compared RTV preceding thought probe reports of being focused and mind wandering. A linear mixed effects model revealed higher RTV prior to mind wandering reports relative to focused reports (See Methods and Fig. 1B for details; *mean ± SD RTV: mind wandering* = 97.94 ± 57.02 vs. *focused* = 67.54 ± 47.11; *t-value*: 2.054, *p=* 0.0403), replicating previous observations (30).

We then tested whether SWRs were differently distributed across attentional states. SWRs were detected automatically using a previously validated approach (3; see Methods). Field potentials were first re-referenced to a bipolar montage (Fig. 2A, top trace), then bandpass filtered between 80–120 Hz (Fig. 2A, middle trace), a frequency range defined *a priori* based on human SWR literature (1,3, 31–36). Candidate ripples were identified as transient high-frequency events exceeding four standard deviations above the mean signal amplitude and lasting at least three oscillatory cycles (Fig. 2A, bottom trace). An averaged SWR waveform from a representative channel is shown in Figure 2B, together with its corresponding time–frequency spectrogram (Fig. 2C). To ensure specificity, interictal discharges were detected separately by thresholding peak-to-trough differences (see Methods) and excluded from all subsequent analyses.

**Figure 2:**
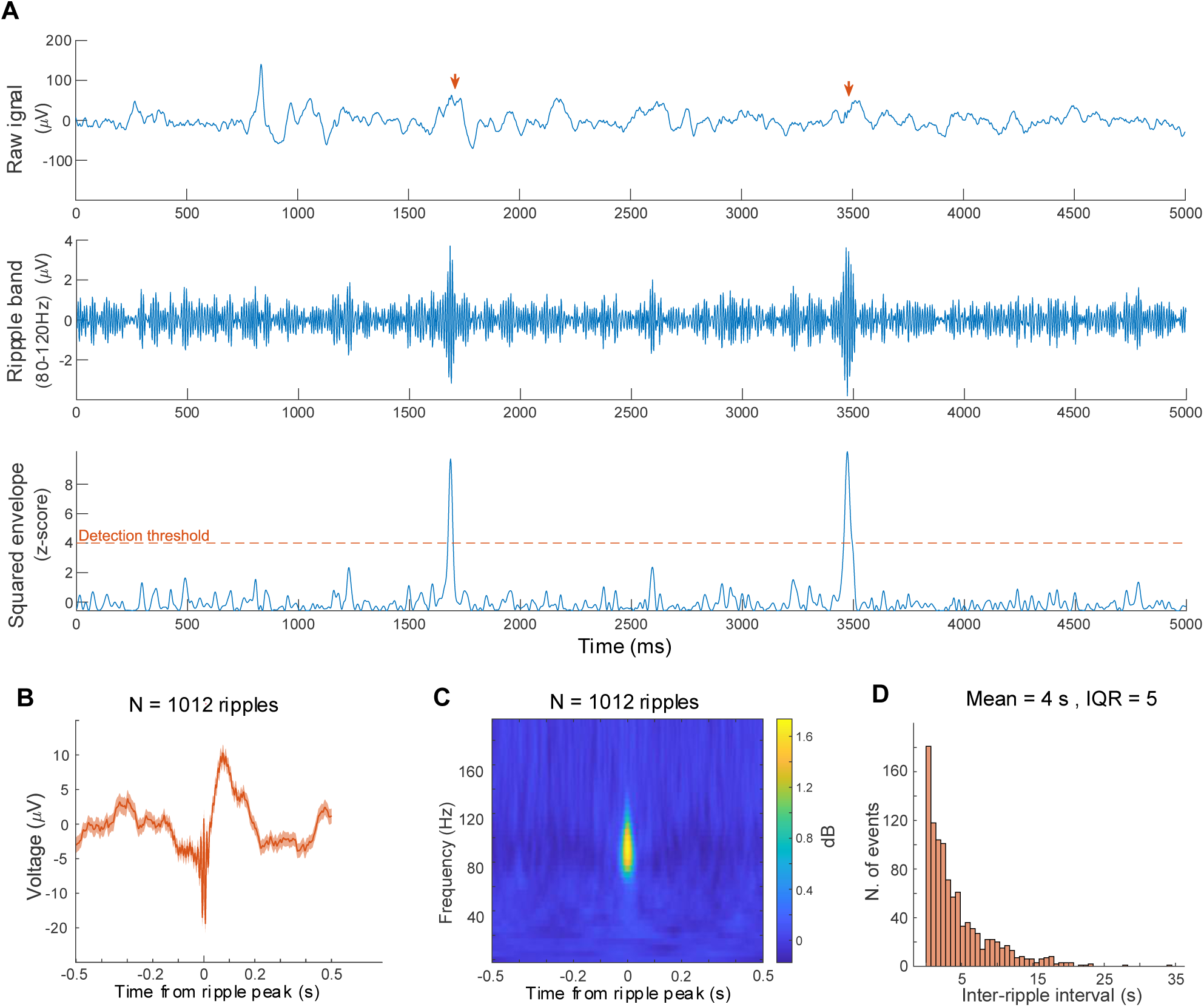
Detection and description of SWRs. **(A)** An overview of SWR detection procedure. Plots present an example of 5 s long snippet of the hippocampal recording. The top plot shows the raw time-course of hippocampal data with two candidate SWR events marked by arrows. The middle panel represents the signal filtered in the SWR range with the same events. The bottom panel shows the envelope of the signal with threshold crossing events as ripple candidates. Horizontal dashed line marks the detection threshold criterion of at least one peak exceeding four standard deviations. **(B)** The field potential locked to the SWR peak. Data come from a representative hippocampal channel. The shading reflects standard error of the mean (SEM). The middle line represents an average of field potential across 1012 SWRs. **(C)** Time-frequency representation of power average of the same 1012 field potentials locked to the SWR peak. Here we notice that the power increase is limited to the range between 80 and 120 Hz. **(D)** The inter-event interval distribution of SWR.

To determine whether attention influences hippocampal SWRs, we examined both the occurrence and the rate of SWRs during online and offline attentional states. Unless otherwise stated, differences between conditions were assessed using a Wilcoxon signed rank test. SWRs occurred more frequently during online than offline states *(mean ± SD:* 233.72 ± 131.83 vs. 214.54 ± 120.47 SWRs; *p* < 0.0001, *z =* 6.27; *n =* 105 channels, *N =* 17 patients; Fig. 3A left panel). Although the overall durations of online and offline states were comparable (see above), subtle timing differences could influence SWR counts. To address this possibility, we computed SWR rates by normalizing SWR count by interval duration. This analysis confirmed significantly higher SWR rate during periods of high attention (*mean ± SD:* 0.2873 ± 0.0830 Hz vs. 0.2580 ± 0.0604 Hz; *p* < 0.0001, *z* = 6.96; Fig. 3A right panel). Control analyses using linear mixed-effects models, which treated attentional state as a fixed effect and participant as a random effect to account for potential clustering, yielded the same pattern of results (see Methods and Supplementary Materials, Table S1). Importantly, both SWR count (*mean ± SD:* 198.26 ± 126.27 vs 179.34 ± 114.99; *p* < 0.001, *z* = 4.39; Fig. 3C left panel) and SWR rate (*mean ± SD:* 0.2758 ± 0.1082 vs 0.2421 ± 0.0810; *p* < 0.001, *z* = 4.81; Fig. 3C right panel) remained significantly higher during online states, after excluding all channels localized in the pathological hemisphere, as identified through the presurgical diagnostics (*n* = 46 channels, *N =* 14 patients; see Methods and Supplementary Materials, Table. S3).

**Figure 3:**
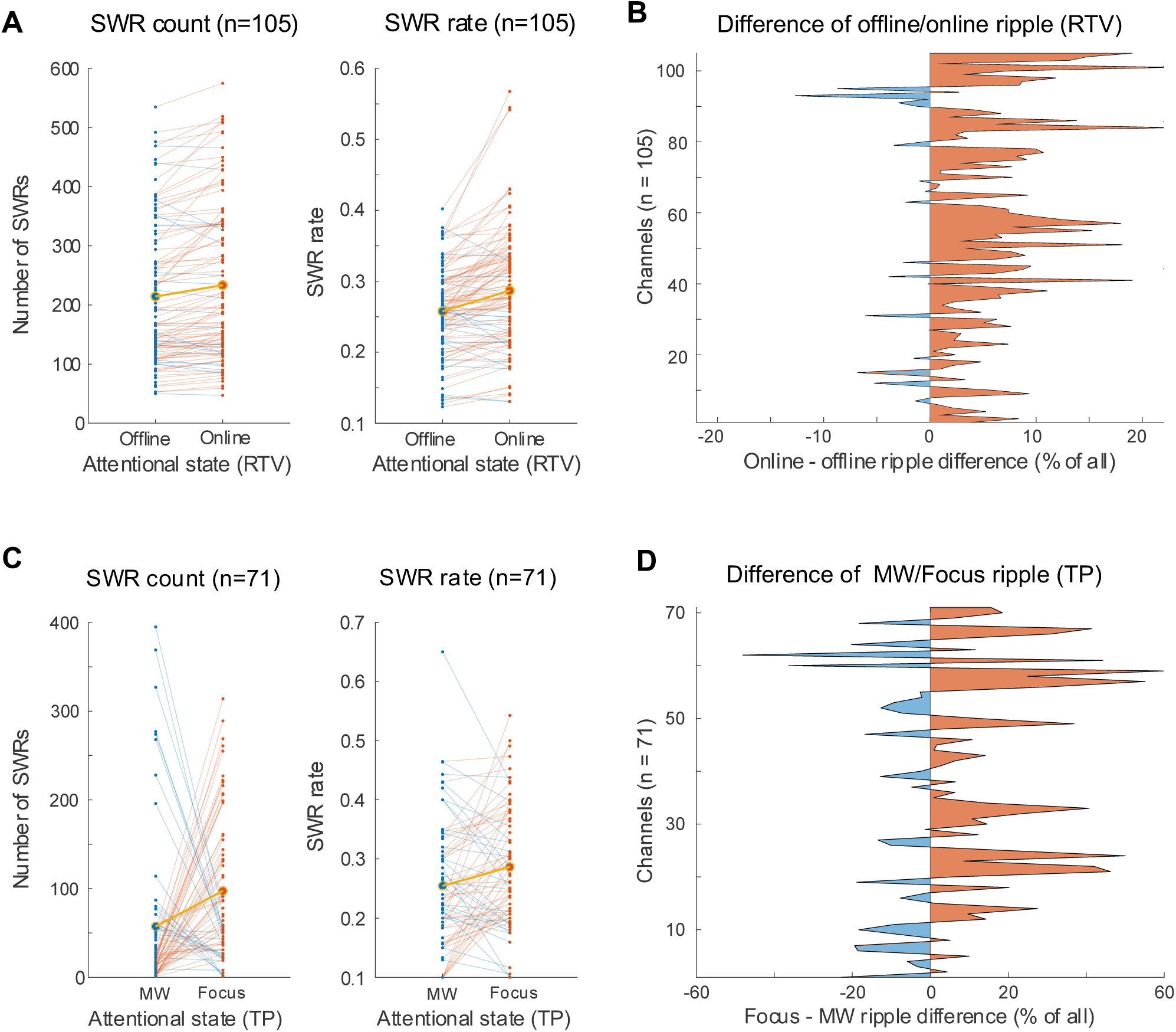
Attention modulation of the SWRs appearance. **(A)** The number (left panel) and rate (right panel) of SWRs during offline and online states. Each line presents a single channel. The mean is marked by large dots and thick orange lines. Each line is color-coded to present the direction of results. If a given channel shows more SWRs during the online state it is marked orange. Otherwise it is marked blue. The SWR rate was computed by dividing the total number of ripples by the total time a participant spent in online or offline state (see Methods). **(B)** Distribution of SWR count across all channels (n = 105), expressed as the normalized difference between online and offline states. For each channel, we computed the difference in SWR count between conditions and divided it by the total SWR count across both conditions. Positive values (orange) indicate a bias towards more ripples during online states; negative values (blue) reflect more ripples during offline states. X-axis presents the magnitude of the difference (in %); y-axis shows individual electrodes. **(C)** The same analysis as in (A), but computed in the 10-second interval preceding each thought probe. Total SWR count (left panel) and SWR rate (right panel). The SWR rate was computed by dividing the total number of ripples by the number of focused or mind wandering responses (see Methods). **(D)** The SWR distribution based on subjective attentional state (Focus vs. MW), expressed as the normalized difference between focused and mind wandering responses. For each channel, we computed the difference in SWR count between thought probes conditions and divided it by the total SWR count across both conditions. Due to some participants providing responses only in one category in a given condition, the total number of included channels was reduced to 71 (see Methods).

Next, we asked whether subjective, introspective reports of attentional state based on thought probes revealed a similar modulation of SWRs. Specifically, we compared SWR rates in the 10 seconds preceding thought probe reports of focused attention versus mind wandering. This window was chosen to capture neural dynamics proximal to introspective judgments while minimizing contamination from post-probe events, as it corresponds to half of the minimum interval between thought probes (see Methods). To ensure the validity of these comparisons, we only included participants who reported experiencing both attentional states. Individuals who consistently reported being either focused or mind wandering were excluded, as such patterns likely reflects a misunderstanding of the introspective task. SWRs were more frequent prior to reports of focused attention (*mean ± SD*: 0.2867 ± 0.1092 Hz) than prior to reports of mind wandering (*mean ± SD:* 0.2545 ± 0.1018 Hz, *p* = 0.0178, *z* = 2.37; *n* = 71 channels, *N* = 12). We further reproduced this observation when excluding channels from the pathological hemisphere: SWR rates remained higher preceding focused-attention reports (*mean ± SD*: 0.2667 ± 0.1212 Hz) compared to mind wandering reports (*mean ± SD* = 0.2192 ± 0.1108 Hz; *p* = 0.0448, *z* = 2.01; *n* = 27 channels, *N* = 12 participants).

Taken together, these results show that hippocampal SWR occurrence is modulated by attentional state, with elevated SWR rates during periods of high sustained attention. Notably, this modulation was evident across both behavioral indices of attention (i.e., reaction time variability) and phenomenological reports (i.e., introspective state reports). These findings suggest that SWRs, typically associated with offline memory processes, are also shaped by moment-to-moment fluctuations in attention, supporting a broader role in processing ongoing experience during wakefulness.

### Attention state modulates SWR-locked phase coherence

Previous studies have shown that low-frequency hippocampal oscillations are influenced by overt, attention (24,37–41), and working memory (41–44). Building on this work, we asked whether attentional state also modulates the temporal coordination between SWRs and ongoing low-frequency neural oscillations. To ensure that observed effects reflected genuine oscillatory structure rather than changes in the aperiodic (1/f) background, we first characterized the spectral profile of hippocampal activity using the FOOOF (Fitting Oscillations and One-Over-F; Fig. 4A; see Methods). This analyses identified three frequency ranges: 3 Hz, 7-11 Hz and 13-19 Hz, consistent with previously reports in the human hippocampus (24,36–39, 40–43).

**Figure 4:**
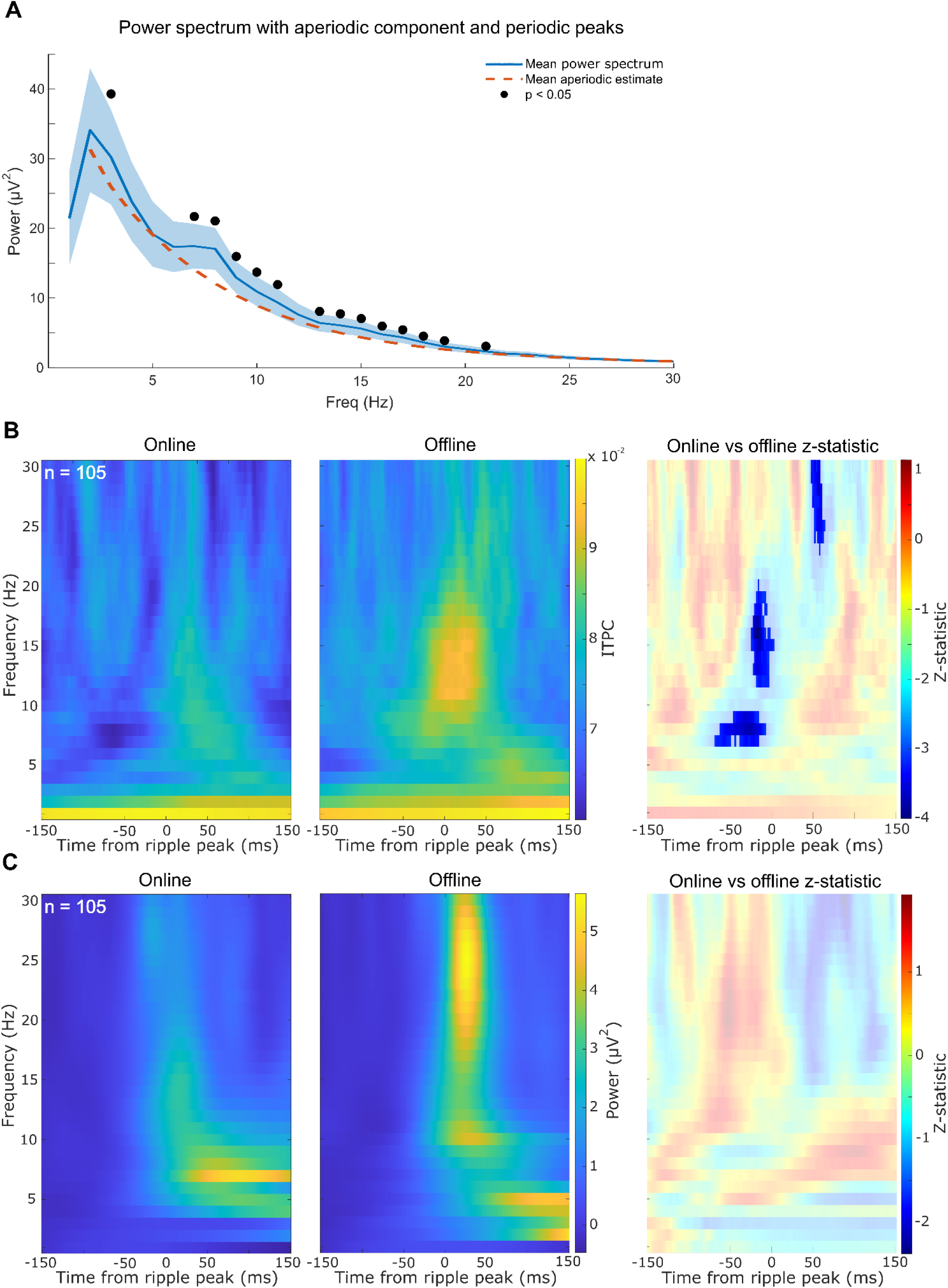
Temporal alignment of SWRs to low-frequency oscillations across attentional states. **(A)** Mean power spectrum across channels. Blue line represents mean power spectrum across channels. Shading reflects standard error of the mean (SEM). Orange dashed line represents a mean fitted aperiodic component to each channel. Black dots above the blue line represent a significant differences between estimated aperiodic model and power spectrum for a given frequency. **(B, C)** Color maps show SWR-locked ITPC **(B)** and power **(C)** during online (left) and offline (middle) states respectively, averaged across channels (n = 105). Right panel presents the z-statistic from Wilcoxon signed rank test of difference between online and offline SWR-locked ITPC (B) and power (C) with mask showing significant time-frequency points after Benjamini-Hochberg correction. Negative z-statistic values indicate that SWRs are better phase aligned during the offline state (B). We detected no systematic changes to SWR-locked power between online and offline attentional states (C).

To test whether SWRs align differently to these oscillatory activity depending on attentional state, we compared SWR-locked phase coherence between online and offline attentional periods. This measure quantifies the consistency with which individual SWRs are aligned to a particular phase of hippocampal neural oscillations (Fig. 4B). Offline states were associated with significantly stronger SWR-locked phase coherence, particularly in the theta and alpha frequency ranges extending to low beta (7–19 Hz), in the 70 ms leading up to the SWR peak (−69 to 0 ms; *all p* < 0.05, *all z* > 1.96; Benjamini-Hochberg FDR correction, *n* = 105). A second cluster of increased coherence was observed 50–60 ms after the ripple peak in the upper beta band (24–30 Hz; *all p* < 0.05, *all z* > 1.96, corrected; Fig. 4B). Control analyses confirmed that these differences were not explained by differences in SWR count across attentional states (see Methods; Fig. S2). In contrast, SWR-locked power analyses (Fig. 4C) revealed only a transient, increase in 20–30 Hz activity during offline states between 0 and 50 ms, which did not survive correction for multiple comparisons and no other differences in SWR-locked power were observed. These findings indicate that temporal alignment of SWRs to ongoing hippocampal oscillations change with changing attentional state. Specifically, during offline intervals – marked by reduced attentional engagement – SWRs exhibited stronger phase locking to low-frequency rhythms, showing enhanced rhythmic coordination under disengaged conditions.

In summary, these results support a model in which attentional state dynamically modulates SWR expression: focused attention associates with increased SWR occurrence, potentially supporting online encoding, whereas disengaged states enhance more precise phase locking of SWRs, potentially reflecting endogenous processes such as memory consolidation. The stronger temporal alignment of SWRs with low-frequency oscillations during offline states may facilitate large-scale network coordination, enabling efficient information transfer or systems-level consolidation across distributed cortical circuits.

## Discussion

We found that both the rate and temporal organization of hippocampal sharp-wave ripples vary systematically with fluctuations in sustained attention. By analysing hippocampal field potentials from presurgical epilepsy patients performing a variant of sustained attention task (i.e., SART), we tracked neural activity across naturally occurring shifts in attentional state, identified using both behavioral measures (reaction time variability) and subjective reports (thought probes). Although SWRs are typically associated with memory consolidation during rest, they occurred more frequently during periods of heightened attentional engagement, even in the absence of explicit memory task. This increase in ripple occurrence was consistent across both behavioral and subjective indicators of attention. Conversely, SWR-locked phase coherence in theta, alpha, and low-beta frequencies was stronger during attention lapses, reflecting enhanced temporal alignment of SWRs with ongoing low-frequency hippocampal rhythms under reduced cognitive engagement. Together, these findings support models in which attention plays a key role in shaping awake SWR activity.

### An awake function of hippocampal SWRs

Our findings point to a broader cognitive function of hippocampal SWRs. Although SWRs have long been primarily associated with rest and sleep (8, 45), accumulating evidence suggests that these high-frequency events also occur during active, goal-directed behavior including encoding (2,13) and visual search (6,7). Here, we show that both the rate of SWRs and their temporal alignment with ongoing low-frequency hippocampal oscillations fluctuate with changes in attentional state.

Elevated SWR rates during focal attentional states may support encoding by enhancing the processing of external information. Consistent with this view, stimuli accompanied by greater ripple occurrence during encoding – presumably under heightened attentional focus – are more likely to be remembered (2). In contrast, the enhanced phase synchronization of SWRs with low-frequency oscillations during reduced attentional engagement may signal a shift toward internally oriented processing (46, 47). The Default Mode Network (DMN), which underlies internally directed thoughts, is reliably recruited during attentional lapses (48,49), and electrophysiological evidence shows that its large scale connectivity is mediated by phase synchronization (50). Importantly, hippocampal SWRs systematically precede DMN activation (51) in primates, suggesting that their activity is coordinated during attentional lapses. Stronger temporal alignment of SWRs to slow oscillations during disengaged periods may therefore facilitate coordinated interactions across distributed cortical regions, supporting efficient information transfer and systems-level consolidation across distributed cortical regions.

Together, our results support a model in which SWRs – and the neural replay they may index – are dynamically gated by attentional state: focused attention increases SWR rate and may facilitate encoding, whereas reduced attention enhances SWR phase locking to low-frequency oscillations, potentially promoting endogenous, consolidation-related processing. This framework aligns with the view that SWR rate and SWR phase synchronization support two distinct aspects of memory: encoding and consolidation, respectively. Such dual-purpose architecture helps explain why SWRs are observed across both externally focused and internally directed cognitive states. These observations are consistent with emerging proposals that task and attention shape the content and expression of neural replay and associated SWRs (52). These results are also consistent with the view that SWRs during reduced attention states promote consolidation processes (18,53) through temporal coordination with neural oscillations.

### Increased rate of awake SWRs during focused attention

Previous studies have shown that a greater number of hippocampal awake SWRs during encoding is associated with better subsequent recall (2, 54,55). In rodents, SWRs occurring at decision points in a maze are essential for spatial learning (12, 56) and higher ripple rates reduce the time spent choosing a path in a maze (57,58), suggesting that increased SWR rates facilitate both learning and decision-making during active wakefulness. Importantly, SWRs are not restricted to the hippocampus: task-related increases in ripple rate have been observed across multiple cortical regions. For example, in the human temporal cortex, the SWR rate scales with the quality of remembered content (59), and in non-human primate visual cortex ripple rates increase when animals maintain visual attention to a stimulus (9).

Our finding that awake SWR rates during periods of heightened attentional engagement aligns with broader literature linking ripple activity to memory formation. Together, these results bridge work across attention, memory, and systems neuroscience, supporting a model in which attention dynamically modulates SWR occurrence. By increasing the likelihood of SWRs, focused attention may enhance the encoding of sensory information through rapid reinstatement mechanisms – similar to those observed during post-encoding periods predictive of subsequent memory performance (60–62). Because SWRs reflect coordinated activation of hippocampal cell assemblies, elevated ripple rates during high attention may signal selective recruitment of neuronal ensembles for encoding. In this framework, attention gates hippocampal reinstatement processes by modulating SWR rate. Beyond encoding, increased SWR rates have also been observed immediately prior to successful recollection (3, 54, 56), potentially reflecting attentional orienting toward internal memory representations (63–67).

Iwata et al. (20) reported higher SWR rates during five-minute windows preceding self-reported internally oriented thoughts. These vivid, imagination-based thoughts differ from the externally directed attentional focus examined here. Importantly, their methodological approach differed from ours in two key respects: (i) SWRs were averaged over long timescales; and (ii) participants were not engaged in an explicit task. In the absence of task demands, ripple activity may reflect quiescent, internally focused states rather than the transient attentional fluctuations captures in our design. Because, attention can be directed both externally and internally, heightened attentional states - and the associated increase in SWR rates - may occur in both contexts depending on cognitive goals. Relatedly, ripple rates increase during brief rest periods interleaved with motor learning trials and predict subsequent performance improvements (68). These moments likely reflect resting states rather than task-driven attentional modulation.

Hippocampal SWRs are also implicated in metabolic control: increases in ripple rate precede reductions in peripheral glucose concentration (69), and this relation is reciprocal. Post-prandial glucose elevation increase SWR occurrence proportionally to caloric load, suggesting that nutrient availability up-regulates hippocampal ripple dynamics, which in turn contribute to restoring glucose homeostasis (70). Transient glucose increases also improve attentional performance (71–73). These findings suggest that glucose availability promotes both SWR generation and attentional engagement, while SWRs themselves feedback to regulate glucose, positioning ripples at a bidirectional interface between metabolic state and cognition.

Extending prior work, we show that even during active task performance, SWR rates track moment-to-moment fluctuations in focal attention. Our results suggest that hippocampal awake SWRs are modulated by attentional state, with elevated rates during heightened attention. At the same time, SWRs also occurred during attentional lapses which, although less frequent, exhibited distinct neurophysiological signatures – suggesting complementary roles for awake SWRs across attentional states.

### The temporal coordination of awake SWRs

As noted above, awake SWRs occurring during attentional lapses exhibit stronger phase synchrony to ongoing hippocampal oscillations. Phase-based temporal coordination has been shown to facilitate memory consolidation (47), suggesting that tighter temporal alignment of SWRs during lapses may support systems-level consolidation. According to the opportunistic consolidation framework (53), the brain exploits brief periods of reduced external engagement to reactivate and stabilize recent experiences through coordinated replay (13,74). Replay propagates across large-scale hippocampo-cortical networks via phase synchronization, a mechanism thought to support synaptic plasticity and long-range information transfer (75). Our findings align with this view by showing that SWRs during attentional lapses become more temporally aligned with low-frequency rhythms.

Prior work during sleep has shown that SWRs couple with cortical spindles and slow oscillations in humans (1, 76) and rodents (77). This coupling predicts subsequent memory performance (78,79) and synchronizes distributed hippocampo-cortical assemblies (32,80). Awake SWRs also align with cortical ripples and high-frequency bursts in anterior temporal and prefrontal regions (2, 3, 54, 75), covarying with cortical theta power (54) and lock to the phase of slow oscillations (59). The increased phase-locking we observe during attentional lapses may therefore contribute to memory consolidation during wakefulness, although future work must determine whether its magnitude predicts behavioral memory outcomes.

Although sleep-associated consolidation is often linked to very slow oscillations (<1 Hz), evidence from awake states suggests that higher-frequency rhythms particularly theta and lower alpha (3–10 Hz) also support memory. Stronger hippocampal-cortical theta coupling during reactivation predicts improved relational memory (81), increases during cued recall (82,83) and scales with retrieval performance (84). Electrical stimulation of the temporal cortex at specific hippocampal theta phases strengthens hippocampo–cortical coupling (85). Optogenetic studies show that theta phase–locked stimulation enhances spatial recall, likely by facilitating pattern reinstatement (83,86,87). While most work focuses on theta, our previous work reveal additional synchronization in the alpha and beta ranges. Theta and alpha activity often covary in the hippocampus (41, 44, 88, 89), reflecting overlapping mechanisms of large-scale coordination.

While phase synchronization during attentional lapses may be important for supporting consolidation during wakefulness (53,90), SWR-mediated synchronization could also contribute to encoding during focused attention. However, during attentive, stimulus-driven states, strong evoked cortical responses may mask or disrupt endogenous SWR-related phase synchronization, making such effects more difficult to detect (see also 24).

More broadly, although consolidation during sleep is classically linked to very slow frequency oscillations (<1 Hz), evidence from awake studies demonstrate that memory performance can benefit from coordination within higher-frequency rhythms, particularly in the theta and alpha ranges.

Together, our findings point to a flexible mechanism: during lapses of externally directed attention, SWRs become more temporally aligned with slower rhythms (i.e., theta/alpha range), enabling integration across hippocampo-cortical networks; during focused attention, SWRs may instead support encoding related computations. Thus, awake SWRs appear to dynamically switch roles – facilitating encoding when attention is externally focused, and contributing to consolidation when attention drifts or lapses.

### Conclusions

Our findings show that fluctuations in sustained attention dynamically shape both the rate and temporal coordination of awake hippocampal SWRs in humans. Elevated ripple rates during focused attention suggest an active role of wake SWRs in encoding, whereas stronger phase coupling to low-frequency oscillations during attentional lapses points to engagement in systems-level consolidation. More broadly, these results support a model in which awake SWRs flexibly adjust to cognitive state – bridging externally directed attention and internally driven processing. By linking moment-to-moment attentional fluctuations to ripple dynamics, our work provides a new insight into how the human hippocampus orchestrates the balance between encoding and consolidation during active wakefulness.

## Methods

### Patients

Intracranial electroencephalography (iEEG) of the mediotemporal lobe (MTL) region together with reaction time (RT) measurements were collected from 18 patients with refractory epilepsy at the Clinic and Polyclinic for Epilepsy at the University Hospital Bonn (mean age 41.42 years; range 20-60). Patients performed a variant of the standard sustained attention to response task (SART) while implanted with depth electrodes for diagnostic purposes. The study was approved by the local ethics committee of the University Hospital Bonn. Out of 18 subjects,17 were included in the analysis. One recording was excluded due to technical failures (the behavioral and electrophysiological time series did not correspond to one another). A Matlab Psychtoolbox (Mathworks Inc., 2022; Brainard, 1996) was used to run the experimental task. Timestamps for presentation of each digit were digitally imprinted on the EEG recording and saved onto the computer used to run the task. Timestamps from behavioral experiment and EEG recording were later manually aligned for each participant, to ensure the correct correspondence.

### Task

Patients performed a variant of the sustained attention to response task (SART). They were presented with a random sequence of digits ranging from 0 to 9 with 2 s inter-digit interval. They were instructed to press the space bar in all cases except when the number ‘3’ was presented, in which case patients were asked to abstain from a response (speed and accuracy were emphasized). After pressing the button, the next digit was displayed after a 2s interval. If no response was made, the procedure re-commenced after 1 s (plus the 2 s interval period). On average patients performed 861.95 trials (*mean* = 861.95, *range* 440; 1440) and took approximately an hour to complete the experimental paradigm *(averaged time* = 51 minutes and *range* from 20 min to 1 hour 30 minutes). Data from responses to the digit presentation sequence sampled reaction time and accuracy. Additionally, from time to time the sequence of digits was interrupted by intermittent thought probes assessing momentary attentional state of the patient (inter-question interval: 20-30 seconds). Patients were asked: “Was your attention focused on the task immediately before the question has been shown on the screen?”. The number of presented probes totalled 120, but the final analysis differed for each patient, due to manual co-registration of behavioral and electrophysiological data. In addition not all patients were able to complete the entire experimental session (*mean* = 77.89, *range* 39 - 120).

### Electrophysiology data acquisition and preprocessing

iEEG data were amplified and recorded using the Neuralynx ATLAS system with bandpass filter between 0.1 and 9000 Hz, and digitized at a rate of 32 kHz. We selected hippocampal electrodes using a standard two-step procedure (i.e., based on a combination of anatomical and physiological criterion). Initially, we identified contacts that were targeting the hippocampus. This step was done by visually inspecting individual implantation schemes for each patient. Subsequently, we analysed the P3 ERP component and then selected channels that were only localized to the hippocampus and that showed a clear MTL-P300 potential evoked by the onset of the digit presentation. The limbic P3 ERP component or the MTL-P300 is generated locally and is elicited by attended stimuli (91), and is an established marker used to identify the hippocampal response (92).

iEEG data underwent minimal preprocessing, Initially, data were resampled to 500 Hz and then residual line noise (at 50 Hz and its harmonics (100, 150 and 200 Hz)) was removed using band-stop filtering. This was achieved by applying a fourth order Butterworth filter with band-stop of +/-2 Hz to account for any variability in line noise across time. Following de-noising, the data were re-referenced to a bipolar montage. Epochs of data containing artefacts were automatically detected using a three-step procedure. First, to detect large abnormalities we rejected 500 ms long segments with amplitudes exceeding 3rd quantile plus 2.3 times of its interquartile range (I R), as adapted from Vaz et al., (3). Second, we removed timestamps where data points exceeded five standard deviations from the mean of the time series (filtered at 250 Hz), due to high frequency noise. Finally, we computed the differential time series (generated with the use of the Matlab function *diff* and thresholded it by five standard deviations, to further account for artefacts. Timestamps of artefacts detected at each stage of processing were marked and saved for later removal.

### Inter-ictal epilepsy discharges removal

The inter-ictal epilepsy discharges (IED) are large amplitude electrophysiological events often observed in patients with epilepsy. To prevent IEDs from influencing our results, we automatically detected and removed them using a standard approach (93). For each iEEG channel, we z-scored the time series across the entire time course. Afterwards, troughs that exceeded four standard deviations from the mean with a duration of 50 ms or less, were identified. If a positive peak occurred within 100 ms after the trough and exceeded four standard deviations of the mean, the whole complex was labelled as an IED candidate. The difference between matched peaks and troughs had to exceed 12 standard deviations to qualify for removal. Timepoints marked as IED were saved and used in subsequent SWR detection. To further ensure that our results were not influenced by IEDs, each SWR candidate was manually inspected to ensure no overlap with any IED event that may not have been previously found by the detection algorithm.

### Sharp Wave Ripple detection

We detected sharp wave ripples (SWRs) using a standard six step algorithm (3). First, epochs within the 80-120 Hz frequency range envelope that exceeded two standard deviations above the mean of the whole time series were identified as potential SWR candidates. Then, those segments that did not contain at least one peak greater than four standard deviations above the mean were excluded. Events closer than 15 ms from each other were concatenated. Candidate events were selected only if they were shorter than 100 ms. Later, via the *findpeaks* Matlab function, only those periods that had at least three peaks were selected, to account for the oscillatory nature of SWRs, and estimating at least three cycles. Finally, the ripple candidates that coincided with epochs containing artefacts or interictal periods were removed. Channels that had very few SWRs (below 50), or did not show a clear SWR when averaging for ripple events, were discarded. This procedure ensured that our analysis was conducted using genuine SWR events. Following the electrode selection process and removal of patient data that did not meet these criteria, 105 channels across 17 patients (mean channels per patient = 6.2) were retained.

### Phase coherence analysis

We computed inter-ripple phase coherence separately for each channel to investigate differences in SWR-locked phase coherence. Each SWR event was classified as either online or offline based on its timestamp alignment with reaction time variability clusters defining the two conditions. For each SWR, we identified the maximum peak (including negative peaks) and extracted a 400 ms time window around it. We used a wavelet decomposition via the FieldTrip function *ft_specest_wavelet* (94), with a cycle of three wavelets spanning frequencies from 1 to 30 Hz to calculate time-frequency representations of signals around SWRs. To minimize edge artefacts, wavelet decomposition was performed over a 5 s period centered on the peak, and the resulting data were cropped to 300 ms. The phase angle of every time frequency point was extracted and used to calculate phase coherence (see Formula 1) across all SWR events for each channel. Differences in phase coherence between online and offline conditions were tested using two-sided Wilcoxon signed-rank tests. To account for multiple comparisons, the Benjamini-Hochberg procedure was applied.

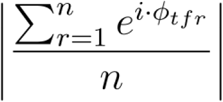

#### Formula 1: Inter-trial Phase Coherence Index

*N is the number of SWRs in condition, while ϕ_tfr_ represents the phase angle of time frequency complex representation*.

#### Phase coherence – control analyses

As phase coherence is sensitive to the number of trials, we conducted control analyses to ensure our results do not depend on the number of SWRs included in the analyses. Initially, this was controlled by randomly downsampling trials. For each channel, we calculated phase coherence, randomly downsampling the number of SWRs to equalize event counts between conditions. Subsequently, we recomputed phase coherence for both conditions (i.e., online and offline) for each downsampled set using the methods described above. This procedure was repeated 1000 times. For each downsampled iteration, we conducted a two-sided Wilcoxon signed-rank test between online and offline conditions, on median ITPC in clustered significant differences of ITPC when controlling for multiple comparisons (see above). This resulted in a distribution of z-statistics from the Wilcoxon test, each corresponding to one of three clusters. (*Cluster 1*: -60 to -14 ms and 7-9Hz; *Cluster 2*: -22 to 0 ms and 11 to 19 Hz; *Cluster 3*: 52 to 60 ms and 24-30 Hz. See Fig. S2 for histograms of z-statistic distributions). The distribution of z-statistics shows that almost all repeated tests resulted in significant statistics (*Cluster 1*: 100 ; *Cluster 2:* 100 *; Cluster 3*: 99.7 ; See Supplementary Materials, Table S4 for details). In sum, control analyses consistently demonstrated significant differences in phase coherence between online and offline conditions in time-frequency regions that exhibited a difference in the initial analysis. This indicates that results showing higher phase coherence in the offline condition do not depend on the number of SWRs included in the analysis (Fig 6).

### Peri-SWR power spectrum and aperiodic modeling

To ensure that the observed ITPC effects reflected locking of SWRs to genuine oscillatory activity rather than differences in the aperiodic (1/f) background, we computed peri-SWR power spectra time-locked to the SWR peak within a ±500 ms window. Power spectra were estimated from 1 to 30 Hz in 1-Hz steps for each channel. The aperiodic component of each spectrum was then modeled using the FOOOF (Fitting Oscillations and One-Over-F) MATLAB wrapper, allowing channel-wise separation of periodic and aperiodic spectral structure. The *knee* parameter was enabled to accurately model spectral flattening in log–log space. For each frequency, the mean empirical power spectrum was compared with the mean FOOOF-estimated aperiodic component across channels using paired *t*-tests. Statistical significance was evaluated using the Benjamini–Hochberg false discovery rate (FDR) correction to account for multiple comparisons across frequency spectrum. This procedure revealed significant clusters of oscillatory power exceeding the aperiodic estimate at 7-11 Hz and 13-19 Hz, broadly overlapping with the frequency ranges exhibiting significant ITPC effects (see Supplementary Materials, Table S4)

### Analyses of behavioral data

To detect periods of offline and online states we used a standard measure of reaction time variability (RTV, 95,96). The reaction time variability time series was calculated using a sliding window of five measurements, moving a step at a time. To estimate variability in reaction times, we calculated standard deviation across these five measurements. The RTV time series was then split, categorizing observations exceeding the median as offline states and those below the median as online states. This measure reflects an established relationship between variance in reaction time responses and sustained attention (95–98). Time points that were no further than 3 s apart and represented the same attentional state, were clustered together and labelled as a single offline or online event. Within each cluster representing online or offline states, SWRs were counted. Additionally, to account for varying durations of each online and offline episode, counts of SWRs were normalized by the total time spent in each state during the whole experimental session. We refer to this measure as the SWR rate. To be certain, that the sliding window length had no effect on our results, we repeated our study applying a sliding window of 10 measurements to calculate instantaneous RTV. These control analyses reproduced previous observations (for SWR count: *mean ± SD*: 230.96 ± 129.55 vs 220.95 ±126.19; *z* = 3.10, *p* = 0.0019; for SWR rate: *mean ± SD:* 0.2825 ± 0.0826 vs 0.2644 ± 0.063; *z:* 4.03, *p* = 0.00006).

### Analyses of subjective thought probes

Additionally, we compared the SWRs rate in time windows preceding responses to thought probes (TP). Here, we calculated the SWR rate in a period equal in length to the half of minimal distance between the current and immediately succeeding thought probe (10 s) for every attentional state inquiry. Patients who showed no response in either of the categories were excluded leaving us with 12 Subjects (71 channels). We computed SWR rate by dividing the total number of SWRs by the summed duration of all 10-second epochs preceding thought-probe reports classified as either focused attention or mind wandering. To assess the relationship between our two measures of attentiveness, we compared RTV values before each thought probe in which participants responded as either mind wandering or focused. Next, we used linear mixed effect random intercept model built as follows: response ∼ condition + thought probe number + (1|patient-id). In this model, the patient identification number was treated as a random effect, while we additionally controlled for time-on-task (i.e., number of the TP).

### Pathological data exclusion

To ensure that the results are not influenced by pathological epileptogenic activity within the hemispheres, we repeated SWR rate analyses with the exclusion of iEEG channels from the hemisphere that contained epileptic seizure onset zones. After the removal of these channels, the data of 14 patients remained in the analyses (two patients had EEG recordings with both hemispheres showing epileptic activity), 46 channels in total. We assessed SWR rate in online and offline states using the same RTV procedure as previously. Subsequently, we also replicated the analyses on the time preceding mind wandering and focused answers for thought probes. Within this case, 27 channels and 12 patients (accounting for skewed focus or MW estimation) remained. Both the difference in SWR count and rate between online and offline states, as well as mind wandering and focused states, still proved significant after excluding all channels from pathological hemispheres.

### Statistics

In the primary analyses presented in the main text, we employed non-parametric rank-based statistics (i.e., Wilcoxon sign-rank tests), treating each electrode contact localized within the hippocampus (n = 105) as an independent unit of observation. Please note that during preprocessing, we implemented steps to mitigate the impact of clustering across channels within individual participants. For example, we re-referenced channels to a bipolar montage which involved subtracting signals from neighboring contacts on each electrode shaft. This procedure increases the independence of signals recorded from contacts within individual patients by enhancing spatial specificity, and minimizing common noise and contributions from distant sources via volume conduction.

To further account for any potential remaining clustering, we conducted control analyses using linear mixed-effects models utilizing the lme4 package in R. A random intercept model was formulated as follows: response ∼ condition + (1|patient-id). In this model, the patient identification number was treated as a random effect. Experimental condition served as a fixed effect, and the magnitude of the signal of interest was the dependent variable. We observed a significant effect of attentional state (offline/online) on SWR count and rate, while controlling for inter-subject variability. (see Supplementary Materials for these results).

In addition, we performed a clustered Wilcoxon test using the R package *clusrank*, to control for subject-level clustering. This analysis also indicates a higher SWR rate and count during online, than in offline periods (see Supplementary Materials for results). Wilcoxon tests were conducted in Matlab and later reproduced in R studio, and the clustered Wilcoxon tests and linear mixed-effects models analyses were both performed using R studio.

## Supporting information

Supplementary Informations

## Acknowledgments

This research was funded by a grant from National Science Center of Poland (Sonata 19: UMO-2023/51/D/HS6/02920).

