## Supplementary Informations for "Attention modulates hippocampal sharp-wave ripples in humans"

Michał Domagała *et al.*

**This PDF file includes:**

Figs. S1 to S2
Tables S1 to S4

**Fig. S1.**
Example implantation scheme from a representative patient. Colored contacts indicate electrode sites localized to the hippocampus and included in subsequent analyses.

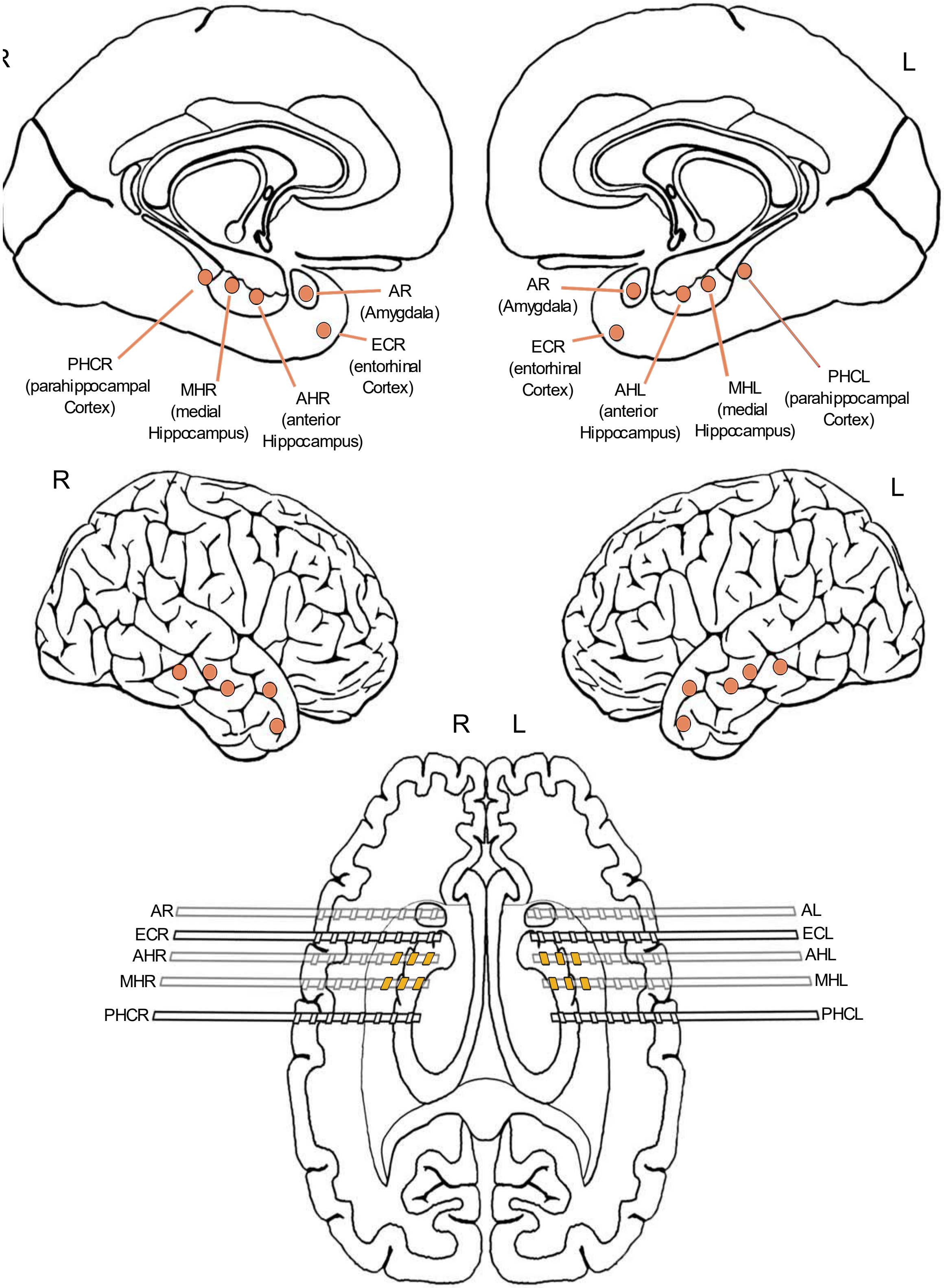

**Fig. S2**

Control analyses for SWR-locked phase coherence. Histograms show the distribution of test statistics from 1000 iterations of downsampling SWR events to match counts between online and offline states. Distributions are plotted for three significant clusters identified in the main analysis: theta (7–9 Hz, –60 to –14 ms), alpha (12–19 Hz, –22 to 0 ms), and beta (24–30 Hz, 52–60 ms). Results confirm that the observed offline > online phase coherence differences are not explained by unequal SWR counts across conditions.

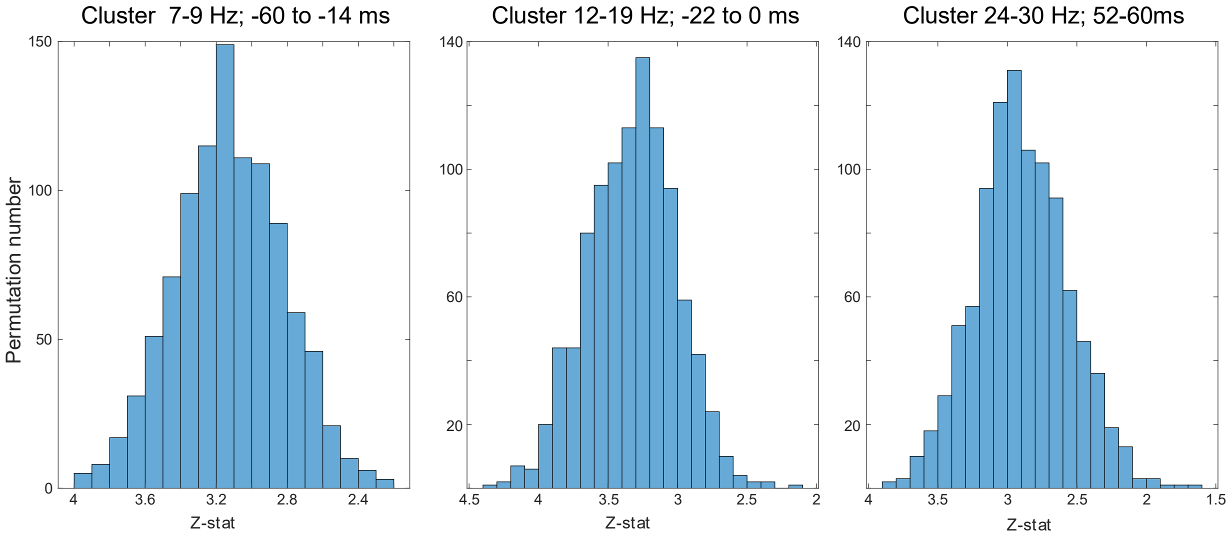

**Table S1.**

Exploration of analysis parameter choices. Results from Wilcoxon signed-rank tests comparing SWR count and SWR rate between attentional states. Analyses were performed using both reaction time variability (RTV; online vs. offline) and thought probe (TP; Focus vs. Mind Wandering) classifications (see Methods). For RTV, our main analysis used a sliding window of 5 responses [RTV (5 responses)], with a control analysis using 10 responses [RTV (10 responses)], yielding comparable results. For TP-based analyses, we compared a 10 s window [TP (10s window)] with a shorter 5 s window [TP (5s window)]. While some differences were observed, we chose the 10 s window as it corresponds to half the minimum inter-probe interval, whereas 5 s may be too brief to reliably capture attentional state. Reported values include Wilcoxon test statistics (z) and corresponding p-values. Overall, results were robust across parameter choices.

| ***Analysis*** | ***Parameter value*** | ***Test name*** | ***Wilcoxon test statistics*** | |
|---|---|---|---|---|
| | | | **z-stat** | **p-value** |
| ***RTV*** | ***5 responses*** | ***SWR count*** | 6.27 | <0.0001*** |
| | | ***SWR rate*** | 6.96 | <0.0001*** |
| ***RTV*** | ***10 responses*** | ***SWR count*** | 3.10 | 0.0019* |
| | | ***SWR rate*** | 4.03 | <0.0001*** |
| ***TP*** | ***10s window*** | ***SWR count*** | 2.91 | 0.0037* |
| | | ***SWR rate*** | 2.37 | 0.0178. |
| ***TP*** | ***5s window*** | ***SWR count*** | 2.90 | 0.0038* |
| | | ***SWR rate*** | 1.37 | 0.1706 |

Significance Codes: 0 ‘***’, 0.001 ‘**’, 0.01 ‘*’, 0.05 ‘.’, 0.1 ‘ ’

**Table S2.**

Results from main and control analysis of SWR frequency in online and offline conditions. Each column corresponds to one of different test for both SWR count and rate, showing both corresponding test statistic and p-value. 'Wilcoxon' column contains values from main analysis. The 'Wilcoxon clustered' column contains results from clustered Wilcoxon analysis to account for inter-subject effect. The 'Mixed models' columns shows both t-statistic and p-values from the linear mixed effect model analysis (online-offline condition ~ SWR measurement + (1|Subject)). These analyses confirm that observed effects were not solely due to inter-subject differences (see Methods)

| ***Test name*** | ***Wilcoxon*** | | ***Wilcoxon clustered*** | | ***Mixed models*** | |
|---|---|---|---|---|---|---|
| | **t-stat** | **p-value** | **z-stat** | **p-value** | **t-stat** | **p** |
| ***SWR Count*** | 6.27 | <0.0001*** | 3.28 | 0.0010* | 2.63 | 0.0091* |
| ***SWR Rate*** | 6.96 | <0.0001*** | 3.48 | 0.0005** | 3.21 | 0.0015* |

Significance Codes: 0 '***', 0.001 '**', 0.01 '*', 0.05 '.', 0.1 ' '

**Table S3:**

Wilcoxon signed-rank results for SWR count and rate after excluding channels within the hemisphere containing epileptogenic zones. Analyses were performed for reaction time variability (RTV; online vs. offline) and thought probe (TP; Focus vs. Mind Wandering) classifications (see Methods). Results confirm that the main effects remained significant after exclusion of pathological channels.

| ***Analysis*** | ***Test name*** | ***Wilcoxon test statistics*** | |
|---|---|---|---|
| | | **z-stat** | **p-value** |
| ***RTV*** | ***SWR count*** | 4.39 | <0.0001*** |
| | ***SWR rate*** | 4.81 | <0.0001*** |
| ***TP*** | ***SWR count*** | 2.70 | 0.0069* |
| | ***SWR rate*** | 2.01 | 0.0448. |

Significance Codes: 0 ‘***’, 0.001 ‘**’, 0.01 ‘*’, 0.05 ‘.’, 0.1 ‘ ’

**Table S4:**

Control analysis of ITPC clusters using downsampling. For each cluster showing significant differences in SWR-locked ITPC between offline and online states, we report its time span (ms) and frequency range (Hz). To control for unequal SWR counts, we repeated the analysis 1000 times, each time randomly downsampling events to match counts between conditions. The table shows the mean and standard deviation of Wilcoxon z-statistics, as well as the percentage of iterations exceeding the critical value ($|z| > 1.96$). Results indicate that all clusters consistently showed stronger ITPC during offline relative to online states.

| ***Cluster number*** | ***Timespan (ms)*** | | ***Freq-span (Hz)*** | | ***Wilcoxon z-stat*** | | |
|---|---|---|---|---|---|---|---|
| | **Start** | **End** | **Start** | **End** | **Mean** | **Std** | **% Correct** |
| **1** | -60 | -14 | 7 | 9 | 3.11** | 0.29 | 100% |
| **2** | -22 | 0 | 12 | 19 | 3.21** | 0.31 | 100% |
| **3** | 52 | 60 | 24 | 30 | 2.91** | 0.32 | 99.7% |

Significance codes: 0 ‘***’, 0.001 ‘**’, 0.01 ‘*’, 0.05 ‘.’, 0.1 ‘ ’.
